# A Comparative Study of Plasma Biomarkers of Neurodegeneration in Rhesus Monkeys (*Macaca mulatta*) and Baboons (*Papio anubis*)

**DOI:** 10.64898/2026.07.02.735676

**Authors:** Michele M. Mulholland, Elizabeth R. Magden, Henrieta Scholtzova, William D. Hopkins

## Abstract

Many nonhuman primate species recapitulate the neuropathological features of sporadic Alzheimer’s disease (AD) to varying degrees. As with humans, the assessment of AD-related pathology in nonhuman primates has historically relied on the use of postmortem brain tissues. *In vivo* alternatives, such as PET imaging tracers and fluid biomarkers, have been developed for use in humans but require further validation in nonhuman primates before replacing postmortem analyses. Here we employed the Nucleic Acid-Linked Immuno-Sandwich Assay (NULISA^TM^) CNS Disease panel to compare age-related changes in plasma biomarkers in two nonhuman primate species (rhesus monkeys and baboons). In addition, we examined whether amyloid and tau biomarkers were associated with brain atrophy, as measured by gray matter volume. We found significant associations between age and multiple biomarkers of neurodegeneration for both species, as well as significant differences in the patterns of these associations between the two species. For the phosphorylated tau measures, though rhesus monkeys had higher values, baboons showed significant and stronger associations with age. By contrast, rhesus monkeys exhibited an earlier age-related decline in Aβ42/Aβ40 ratio than baboons. Finally, in both species, lower Aβ42/Aβ40 ratios were associated with lower gray matter volumes. This is the first systematic comparative study of age-related changes in neurodegeneration biomarkers in two closely related nonhuman primates using comparable age ranges and sample sizes, and the same multiplex assay. Future studies should examine longitudinal changes in these biomarkers as well as validate the plasma findings using cerebral spinal fluid.

## Introduction

Alzheimer’s disease (AD) is a neurodegenerative disorder affecting more than 55 million people worldwide and is expected to rise as life expectancy increases and the number of aged individuals continues to climb (Figures, 2026). Pathologic hallmarks of AD include aggregations of the amyloid-beta (Aβ) protein into extracellular plaques, tau-associated neurofibrillary tangles (NFT), extensive neuronal and synaptic loss, mitochondrial dysfunction, neuroinflammation, and severe cognitive impairment (Montine et al., 2012). In addition, many AD cases involve cerebral amyloid angiopathy (CAA), a condition in which Aβ build up occurs in the brain’s blood vessels (Greenberg & van Veluw, 2024; Jellinger, 2002). Although AD is considered unique to humans, rodent models have played a significant and valuable role in understanding the disease, and they continue to be the primary animal model in therapeutic preclinical trials (Singhaarachchi et al., 2025). However, many AD therapeutics that show promise during preclinical development in rodent models fail to elicit similar effects in humans (Granzotto et al., 2024; Quinn, 2018). For this reason, there is a need for alternative animal models that more closely recapitulate human AD pathology, as well as innovative approaches to facilitate the discovery of more effective treatments and interventions.

There has been a resurgence of interest in developing primate models for preclinical studies of AD and related dementias due to the fact that many species, including prosimians, platyrrhines, catarrhines, and great apes naturally recapitulate neuropathological features of sporadic AD (Arnsten et al., 2021; Barnes et al., 2024; Cramer et al., 2018; Diehl Rodriguez et al., 2024; Edler et al., 2017; Geula et al., 2002; Paspalas et al., 2018; Perez-Cruz & Rodriguez-Callejas, 2023; Perez et al., 2013; Rosen et al., 2008; Schultz, Dehghani, et al., 2000). For example, amyloid plaques, including CAA, are found in the brains of aged individuals in multiple primate species (Elfenbein et al., 2007; Patel et al., 2021; Zeiss et al., 2025). NFTs appear to be less common in nonhuman primates, although they have been reported in some species at extreme old age. (Barnes et al., 2024; Colwell & Emborg, 2025; Edler et al., 2017; Toussaint et al., 2025).

Traditionally, AD-related neuropathology in humans has been identified using postmortem brain tissues (Grothe et al., 2021; Molinuevo et al., 2018; Salvado, Ossenkoppele, et al., 2023; Smirnov et al., 2022). As an alternative, studies have shown that fluid biomarkers of neurodegeneration, including proteins linked to AD core pathology, such as Aβ40, Aβ42, total tau, and phosphorylated tau (e.g., ptau-181, ptau-217, etc.) can be quantified in CSF, plasma, and serum and these measures have been shown to correlate with loss in cognitive functions (Aschenbrenner et al., 2022; Mattsson-Carlgren et al., 2023; Pais, Forlenza, & Diniz, 2023; Pais, Kuo, et al., 2023; Salvado, Larsson, et al., 2023). In addition, AD neuropathology can now be characterized in living patients using PET imaging (Bejanin et al., 2017; Johnson et al., 2016; Soleimani-Meigooni et al., 2020), with several FDA-approved radiotracers available for the detection of AD-related pathology (e.g., fluortaucipir, fluorbetapir). The development and implementation of fluid biomarkers in research on neurodegeneration are important from both clinical and research perspective because they provide a relatively inexpensive and non-invasive means of potentially characterizing acute and longitudinal changes in neuropathology. Moreover, the use of fluid biomarkers provides the opportunity to develop early diagnostic tools and monitor disease progression, particularly in relation to clinical outcomes and responses to different interventions.

The goal of this study was to compare age-related changes in biomarkers of neurodegeneration in two species of nonhuman primates namely baboons and rhesus monkeys. To date, there are only a few studies that have quantified AD biomarkers in either plasma or CSF in relation to age, brain, behavior or neuropathology in nonhuman primates (Beckman et al., 2021; Chen et al., 2018; Latimer et al., 2019; Neal et al., 2025; Phillips et al., 2024; Robertson et al., 2022). Of particular interest was the comparison in fluid biomarker measures between rhesus monkeys (*Macaca mulatta*) and baboons (*Papio anubis*), two closely related catarrhine primate species with comparable lifespan and healthspan (Huber et al., 2025) that often serve as models for research on the comparative biology of aging and neurodegeneration (Barnes et al., 2024; Colwell & Emborg, 2025; Zeiss et al., 2025). Despite their many similarities, these two species appear to differ in the extent of tau pathology and amyloid deposition based on the available published neuropathology data. Notably, Schultz, Hubbard, et al. (2000) reported that 70% of baboons over the age of 20 years exhibit mild to severe tau pathology in the temporal cortex with minimal Aβ deposition and no relationship with Aβ, suggesting they are an excellent model species for studying tauopathy. By contrast, despite the presence of significant amyloid plaques, other catarrhines such as rhesus monkeys and related macaque species exhibit less tau pathology, including NFTs, which appear to occur only in very old individuals (Barnes et al., 2024; Colwell & Emborg, 2025; Paspalas et al., 2018). These findings suggest that there are features of AD pathology that may differ between closely related primate species.

Three limitations in previous comparative studies on biomarkers of neurodegeneration include: (1) the type of biomaterial analyzed can vary across studies (i.e., CSF, plasma, serum), (2) different assays are often used to quantify the biomarkers thereby making direct comparisons between studies difficult, and (3) age ranges and the proportional representation of males and females can vary substantially across reports. Here, we addressed these limitations by analyzing plasma samples in both rhesus monkeys and baboons using the same multiplex assay. Further, we tried to balance the age ranges and standardize the male-to-female ratio between species to enhance the validity of the comparison and minimize these factors as potential confounds. Based on the existing AD neuropathology data, as well as previous reports on fluid biomarkers of neurodegeneration in these two species, we hypothesized that baboons would exhibit higher levels of phosphorylated tau (pTau) compared to rhesus monkeys, whereas the opposite pattern would be found for the Aβ measures. We further hypothesized that the associations between age and pTau measures would be stronger in baboons compared to rhesus monkeys.

Finally, we examined whether amyloid and tau biomarkers were associated with measures of cortical atrophy in the brains of the rhesus monkeys and baboons. In humans, the brain becomes increasingly atrophied in older individuals, as evidenced by declines in brain weight, and gray and white matter volumes, as well as enlargement of the ventricles (Fujita et al., 2023; Pini et al., 2016). The morphological changes in the brain associated with age are hypothesized to be related to decline in microstructural organization including neuronal loss, reduced synaptic density, and demyelination of the cortex. In neurodegenerative disorders, such as AD or fronto-temporal dementia, brain atrophy can be accelerated and particularly severe due to the accumulation of neuropathological changes, such as the buildup of amyloid plaques and neurofibrillary tangles in the cortex (Double et al., 1996; Planche et al., 2021). Like humans, total intracranial, gray and, to a lesser extent, white matter volume loss has also been reported with increasing age in multiple nonhuman primate species (Alexander et al., 2008; Autrey et al., 2014; Darusman et al., 2014; Dash et al., 2023; Didier et al., 2016; Frye et al., 2022; Herndon et al., 1999; Koo et al., 2012; Makris et al., 2007; Phillips & Sherwood, 2012; Sherwood et al., 2011; Westerhausen et al., 2020; Westerhausen & Meguerditchian, 2021; Wisco et al., 2008). With respect to ventricle volume, we identified only two studies testing for associations with age. Both reports examined rhesus monkeys and found significant positive associations between age and the third ventricle volume (Dash et al., 2023; Wisco et al., 2008). Here, we quantified gray matter volumes using magnetic resonance image (MRI) scans in a subset of rhesus monkeys and baboons. We tested for associations between the neuroanatomical and fluid biomarkers measures while controlling for sex and age.

## Methods

### Subjects

There were a total of 137 subjects including 62 rhesus monkeys (*Macaca mulatta*; 53 females, 9 males) and 75 olive baboons (*Papio anubis*; 66 females, 9 males) housed at the Michale E. Keeling Center for Comparative Medicine and Research within UT MD Anderson Cancer Center. Rhesus monkeys ranged between 8 and 29 years of age (*Mean* = 19.65 years, *SE* = 0.72) while the baboons ranged between 9 and 24 years of age (*Mean* = 16.67, *SE* = 0.71). All animals were housed in social groups with environmental enrichment, were fed a diverse and species-appropriate diet, and had *ad libitum* access to water. Rhesus monkeys were housed in sheltered outdoor enclosures, and baboons in either corrals or Primadomes^TM^ with indoor/outdoor access. All procedures complied with federal regulations and guidelines and were approved by the Institutional Animal Care and Use Committee at UT MD Anderson Cancer Center.

### Blood Collection and Analysis

For both species, blood samples were collected at the time of the subject’s annual physical exam or when sedated for other procedures (e.g. brain imaging). The animals were sedated with an intramuscular (IM) injection of 10 mg/kg ketamine (and if sedated for imaging, maintained on 1-3% isoflurane inhalation). Clinical staff collected ≤ 6 ml of blood (EDTA) from a peripheral vein. Following collection, whole blood was centrifuged at 1500g for 10 minutes at room temperature and plasma was aliquoted and transferred to a new tube for storage at -80 degrees C.

The plasma samples were analyzed using the NULISAseq CNS Disease Panel 120 (Next-generation Ultra-sensitive Ligand-based Immunoassay; Alamar Biosciences) run by the MD Anderson-supported <u>O</u>ncology <u>R</u>esearch and Immuno-m<u>ON</u>itoring (ORION) Core. Specifically, the NULISA^TM^ platform uses immunoassay technologies with ultra-high sensitivity and dynamic range for protein biomarkers extracted from biological fluids. NULISAseq is a multiplex assay that employs quantification of proteins relative to a chosen standard and the values are expressed as units of fold change rather than absolute values (i.e., pg/mL). The primary outcome measure from this platform is the NULISA Protein Quantification (NPQ), which is an Alamar-specific relative quantification unit for NULISAseq assays. NPQ is calculated by normalizing NGS read counts by an internal control, normalizing using interplate control samples, rescaling the data, and then performing a log2 transformation. A one unit increase in NPQ represents a doubling or two-fold change in the target protein level. The NULISAseq CNS Disease Panel 120 characterizes neurodegeneration across 120 biomarkers broadly classified into eight categories of CNS function including (1) pathological protein aggregation, (2) synaptic and neuronal network defects, (3) aberrant proteostasis, (4) cytoskeletal abnormalities, (5) altered energy homeostasis, (6) DNA and RNA defects, (7) inflammation, and (8) neuronal cell death. In this study, we were specifically interested in the subset of biomarkers that included amyloid beta (Aβ38, Aβ40, Aβ42), phosphorylated tau (pTau-181, pTau-217 and pTau-231), total tau (MAPT), acetylcholinesterase (ACHE), beta-site APP-cleaving enzyme (BACE1), brain acid-soluble protein (BASP1), CD63, cystatin-C (CST3), glial fibrillary acidic protein (GFAP), insulin-like growth factor-binding 7 (IGFBP7), kallikrein-related peptidase 6 (KLK6), neurofilament light (NfL), presenilin-1 (PSEN1), and secreted frizzled-related protein 1 (SRFP1).

### Magnetic Resonance Imaging

Brain images, using magnetic resonance imaging (MRI), were previously collected from a subset of the subjects, including 32 rhesus monkeys (25 F, 7 M; 8-29 yo) and 72 baboons (63 F, 9 M; 4-24 yo). All images were acquired on a mobile 1.5 Tesla scanner (Phillips Achieva) with either an 8-channel head coil (young males and all female baboons and all rhesus monkeys) or a 6-channel flex body coil (adult male baboons only). All subjects were sedated using ketamine (10 mg/kg IM), intubated, and maintained on 1-3% isoflurane anesthesia until the ∼60-minute scan session was completed. They were placed in the scanner in a supine position, and structural T1-weighted images were acquired (TR: 17 ms; TE: 3.7 ms; flip angle: 13°; number of signals averaged: 10 for rhesus and 8 for baboons; matrix size: 133 x 320; voxel resolution: 0.8 mm). After completing MRI procedures, all subjects were temporarily singly housed adjacent to their home enclosure until fully recovered from the anesthesia, after which they were returned to their social group. Following image acquisition, each MRI scan was aligned in the ACPC axis, skull-stripped, bias corrected and denoised following procedure described in detail elsewhere (Hopkins et al., 2025). Images were then segmented into binary volumes of gray matter, white matter, and cerebral spinal fluid, and the gray matter volume (in mm^3^) was measured using the FMRIB Software Library Tools (FSL; Oxford; https://fsl.fmrib.ox.ac.uk/fsl/docs/).

### Data Analysis

Because NULISA has not been used in previous biomarker studies in nonhuman primates, detection rates for each biomarker within each species are presented in Supplemental Table 1 for completeness and as future comparative reference points. Cross-reactivity across all biomarkers on the panel was approximately 70% for both rhesus monkeys and baboons. That said, there was 100% cross-reactivity for all the biomarkers of interest to this study. Prior to all analyses, within each species, boxplots were created for each biomarker, and any data point falling outside the 99^th^ percentile were removed. This was fewer than 3 individuals per marker within each species. In addition to the NPQ data for the individual markers, we created three ratio measures that have been reported to accurately discriminate between individuals who are or are not at risk for the onset of AD or related dementias (Aschenbrenner et al., 2022; Pais, Forlenza, & Diniz, 2023; Smirnov et al., 2022; Wang et al., 2025). The ratio measures included Aβ42/Aβ40, pTau-217/Aβ42 and pTau-217/total tau. Because the NPQ values were the result of a logarithmic transformation, we subtracted the denominator term from the numerator terms to derive each ratio. All data were analyzed using inferential statistics (i.e., regression, correlation, analysis of variance) with alpha set to *p* < .05.

## Results

### Within Species Associations Between Age and Each Biomarker

We initially tested for linear and quadratic associations between age and each biomarker within species using stepwise multiple regression (see Table 1). Linear and quadratic age were entered into the model and the change in *F*-value was computed to determine whether the quadratic association accounted for a significantly greater proportion of variance in the biomarker measure than the linear model. For the rhesus monkeys, significant quadratic associations were found between age and KLK6 and pTau-217/MAPT while significant linear associations were found for Aβ38, Aβ40, GFAP, NfL, and the Aβ42/Aβ40 ratio. For the baboons, significant quadratic associations were found between age and Aβ42, CST3, pTau-181, Aβ42/Aβ40, and pTau-217/Aβ42. Significant linear associations were found between age and Aβ38, Aβ40, ACHE, BACE1, KLK6, NfL, PSEN1, pTau-217, pTau-231, and SRFP1. Scatterplots with the best fit lines for each biomarker and species are shown in Figure 1A-F.

**Figure 1.**
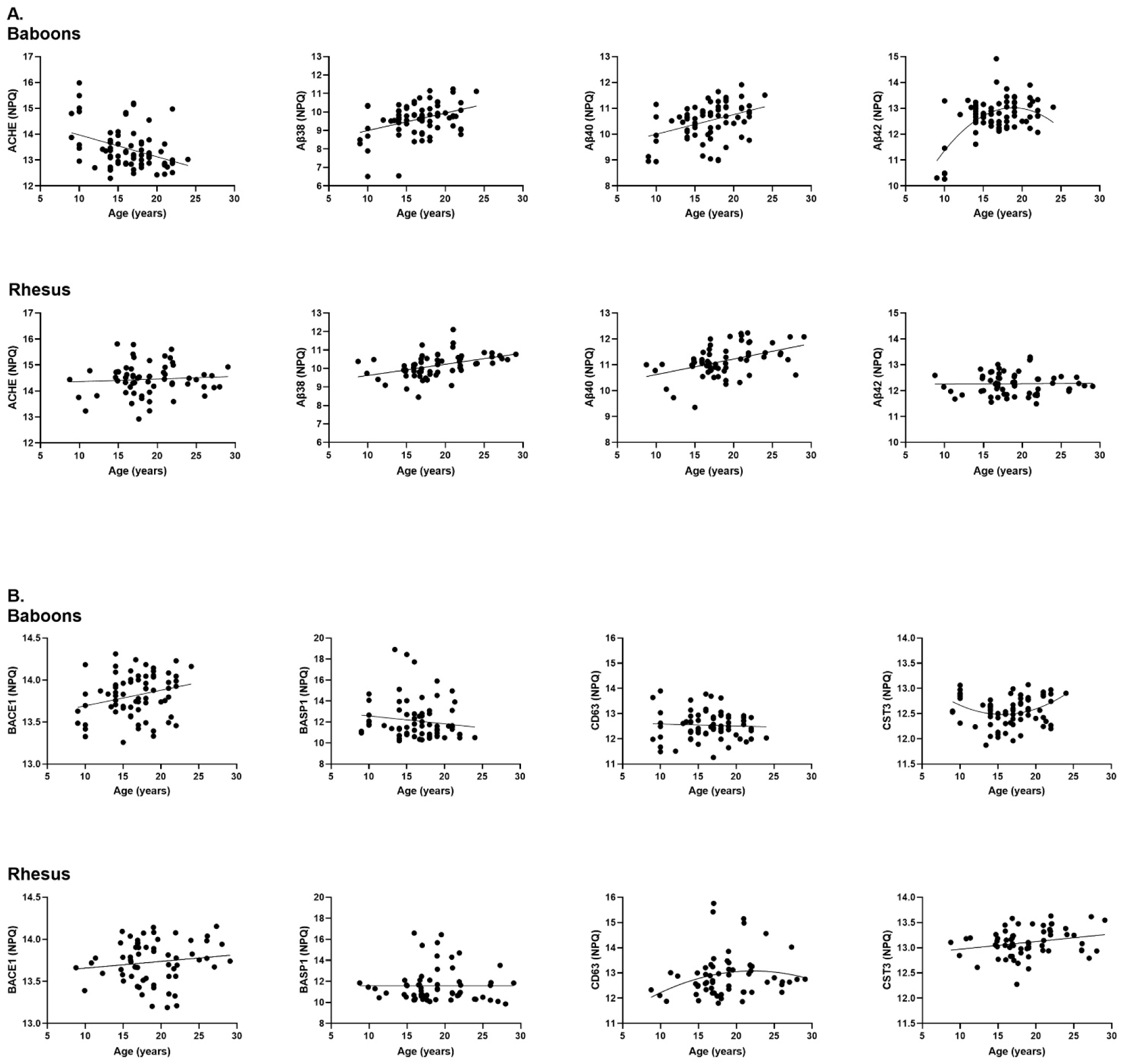
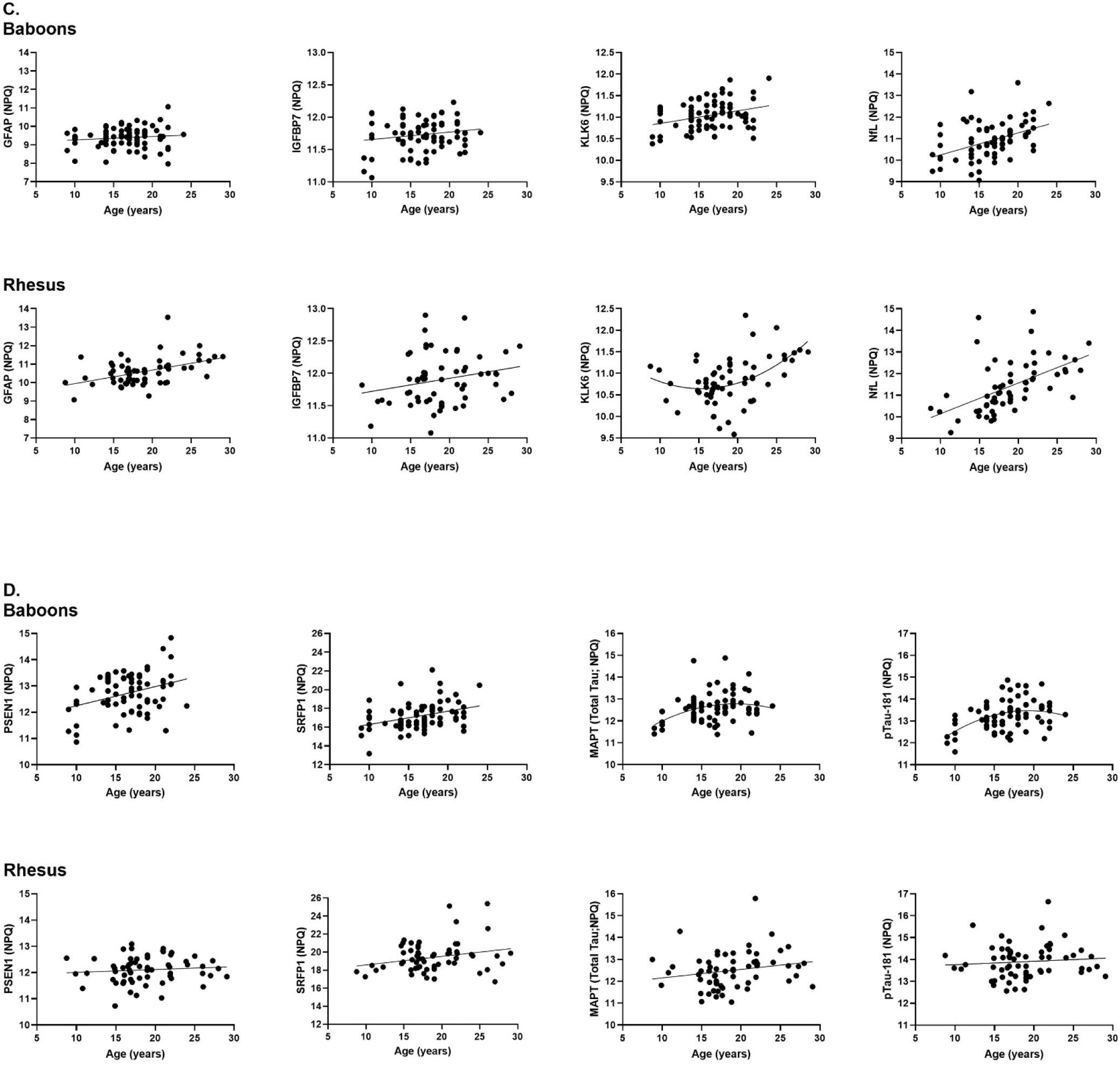
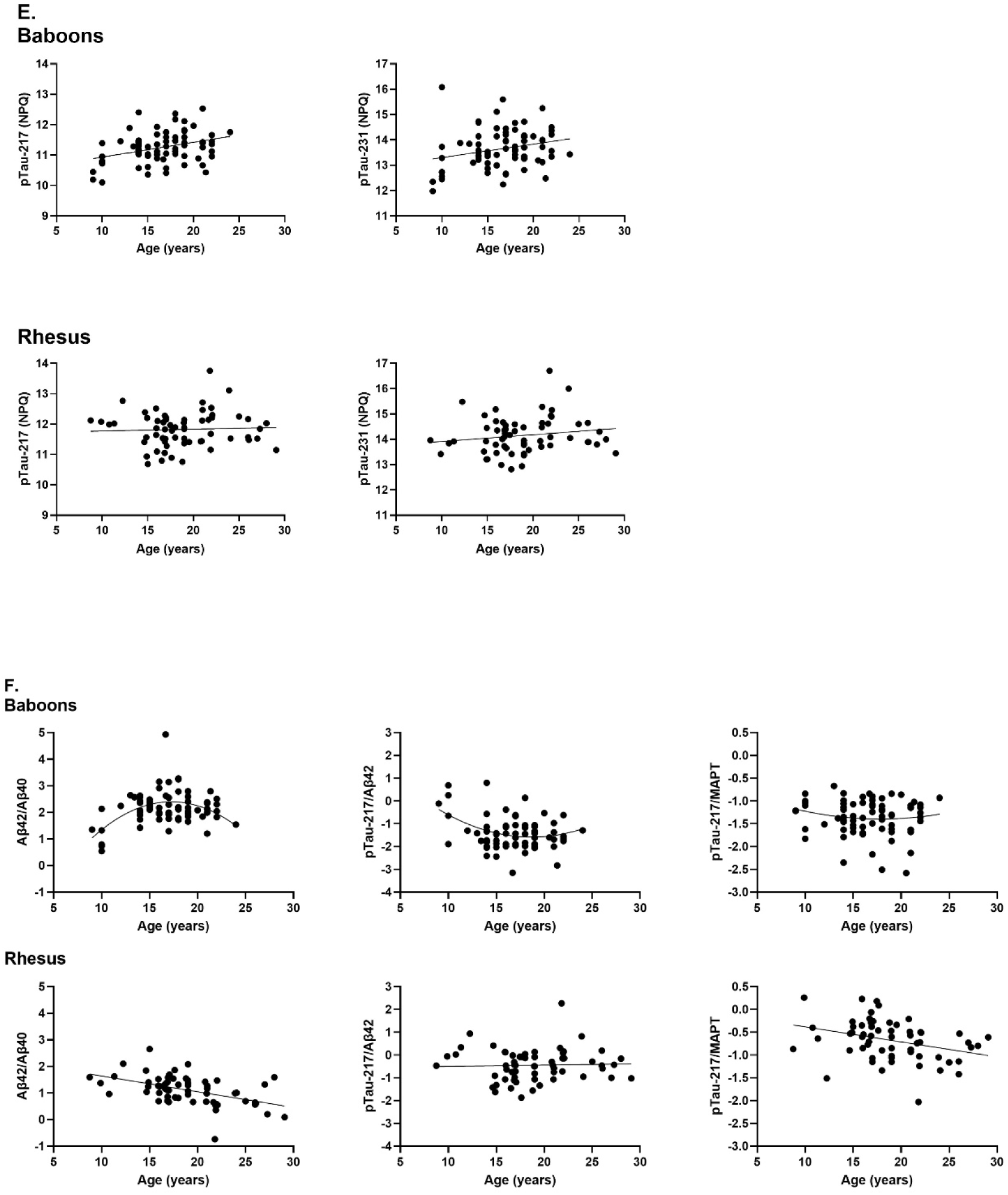
Scatterplots showing the associations between age and each biomarker for both baboons and rhesus monkeys: (A.) ACHE, Aβ38, Aβ40, (B.) BACE1, BASP1, CD63, CST3, (C.) GFAP, IGFBP7, KLK6, NfL, (D.) PSEN1, SRFP1, MAPT, pTau-181, (E.) pTau-217, pTau-231, and (F.) Aβ42/Aβ40, pTau-217/Aβ42, and pTau-217/MAPT.

**Table 1.** Association Between Age and Each Biomarker Within Rhesus Monkey and Baboon Samples.

|  | Linear |  |  | Quadratic |  |  |
| --- | --- | --- | --- | --- | --- | --- |
|  | F-value | r<sup>2</sup> | p | F-value | r<sup>2</sup> | p |
| *Rhesus (n = 62)* |  |  |  |  |  |  |
| Aβ38 | 13.490 | 0.186 | 0.001 | 0.240 | 0.190 | 0.626 |
| Aβ40 | 15.000 | 0.200 | 0.001 | 0.250 | 0.203 | 0.619 |
| Aβ42 | 0.010 | 0.000 | 0.905 | 0.540 | 0.009 | 0.466 |
| ACHE | 0.030 | 0.006 | 0.566 | 0.990 | 0.022 | 0.323 |
| BACE1 | 1.270 | 0.021 | 0.265 | 0.450 | 0.028 | 0.503 |
| BASP1 | 0.000 | 0.000 | 1.000 | 1.170 | 0.020 | 0.283 |
| CD63 | 1.800 | 0.029 | 0.184 | 2.050 | 0.062 | 0.157 |
| CST3 | 3.810 | 0.060 | 0.056 | 0.040 | 0.060 | 0.849 |
| GFAP | 15.480 | 0.205 | 0.001 | 0.640 | 0.214 | 0.428 |
| IGFBP7 | 3.150 | 0.054 | 0.070 | 0.720 | 0.065 | 0.400 |
| KLK6 | 10.750 | 0.152 | 0.002 | 6.300 | 0.234 | 0.015 |
| MAPT | 1.440 | 0.023 | 0.235 | 2.280 | 0.060 | 0.136 |
| NfL | 23.200 | 0.279 | 0.001 | 0.010 | 0.279 | 0.939 |
| PSEN1 | 0.580 | 0.010 | 0.450 | 0.220 | 0.013 | 0.641 |
| pTau-181 | 0.490 | 0.008 | 0.499 | 0.010 | 0.008 | 0.933 |
| pTau-217 | 0.130 | 0.002 | 0.722 | 0.220 | 0.006 | 0.643 |
| pTau-231 | 1.680 | 0.027 | 0.200 | 0.440 | 0.034 | 0.509 |
| SRFP1 | 3.780 | 0.059 | 0.057 | 0.840 | 0.072 | 0.363 |
| Aβ42/Aβ40 | 18.340 | 0.237 | 0.001 | 0.010 | 0.237 | 0.930 |
| pTau-217/Aβ42 | 0.080 | 0.001 | 0.775 | 0.690 | 0.013 | 0.408 |
| pTau-217/MAPT | 1.350 | 0.022 | 0.249 | 6.590 | 0.120 | 0.013 |
| *Baboon (n =75)* |  |  |  |  |  |  |
| Aβ38 | 11.910 | 0.142 | 0.001 | 1.160 | 0.156 | 0.286 |
| Aβ40 | 11.990 | 0.145 | 0.001 | 0.410 | 0.150 | 0.523 |
| Aβ42 | 18.800 | 0.212 | 0.001 | 16.840 | 0.366 | 0.001 |
| ACHE | 12.250 | 0.144 | 0.001 | 2.740 | 0.175 | 0.102 |
| BACE1 | 5.460 | 0.070 | 0.022 | 0.740 | 0.079 | 0.392 |
| BASP1 | 1.610 | 0.022 | 0.209 | 0.810 | 0.032 | 0.371 |
| CD63 | 2.890 | 0.038 | 0.093 | 1.030 | 0.052 | 0.314 |
| CST3 | 0.210 | 0.003 | 0.648 | 8.040 | 0.103 | 0.006 |
| GFAP | 1.050 | 0.014 | 0.309 | 0.440 | 0.020 | 0.510 |
| IGFBP7 | 2.140 | 0.028 | 0.148 | 1.290 | 0.045 | 0.260 |
| KLK6 | 8.250 | 0.102 | 0.005 | 0.480 | 0.107 | 0.493 |
| MAPT | 0.080 | 0.001 | 0.783 | 2.350 | 0.033 | 0.130 |
| NfL | 16.960 | 0.188 | 0.001 | 1.620 | 0.206 | 0.207 |
| PSEN1 | 10.130 | 0.122 | 0.002 | 1.790 | 0.143 | 0.185 |
| pTau-181 | 11.480 | 0.137 | 0.001 | 5.350 | 0.198 | 0.024 |
| pTau-217 | 9.310 | 0.116 | 0.003 | 3.510 | 0.158 | 0.065 |
| pTau-231 | 4.700 | 0.061 | 0.033 | 1.840 | 0.084 | 0.179 |
| SRFP1 | 10.321 | 0.124 | 0.002 | 0.000 | 0.124 | 0.999 |
| Aβ42/Aβ40 | 3.980 | 0.005 | 0.050 | 21.360 | 0.281 | 0.001 |
| pTau-217/Aβ42 | 5.140 | 0.070 | 0.027 | 4.990 | 0.134 | 0.030 |
| pTau-217/MAPT | 0.080 | 0.001 | 0.777 | 0.410 | 0.007 | 0.524 |

Aβ38 – amyloid beta 38, Aβ40 – amyloid beta 40, Aβ42 – amyloid beta 42, ACHE – acetylcholinesterase, BACE1 – beta-site APP-cleaving enzyme, BASP1 - brain acid-soluble protein, CD63, CST3 – cystatin C, GFAP - glial fibrillary acidic protein, IGFBP7 – insulin-like growth factor-binding 7, KLK6 – kallikrein-related peptidase 6, MAPT – total tau, NfL – neurofilament light, PSEN1 – presenilin-1, pTau-181 – phosphorylated tau 181, pTau-217- phosphorylated tau 217, pTau-231 - phosphorylated tau 231, SRFP1 – secreted frizzled-related protein 1.

### Slope Differences Between Species

Here we tested whether the slope in age-related changes in each biomarker differed between species, with sex and species as the between-group factors and age as the covariate. We built an interaction term between species and age into the model with significant F-values indicating a significant difference in slope. The results of these analyses are shown in Table 2. Baboons had significantly greater slopes in change compared to the rhesus monkeys for Aβ42, ACHE, PSEN1, and the pTau-217/Aβ42 ratio. Rhesus monkeys had greater slopes in change for, GFAP, and the Aβ42/Aβ40 ratio. We note here that these analyses tested for differences in the linear changes between species but, as is clear from Table 1, for some markers, quadratic associations account for a greater proportion of variance across ages (e.g. the Aβ42/Aβ40 and pTau-217/Aβ42 ratios).

**Table 2.** Comparisons in Slope of Change in Each Biomarker Between Species.

| *Individual Measures* | Rhesus<br>M (SEM) | Baboon<br>M (SEM) | *F*-value | *p*-value |
| --- | --- | --- | --- | --- |
| Aβ38 | 0.053 (0.016) | 0.094 (0.028) | 1.42 | 0.236 |
| Aβ40 | 0.053 (0.015) | 0.074 (0.022) | 0.66 | 0.417 |
| **Aβ42** | **-0.004 (0.012)** | **0.104 (0.024)** | **17.15** | **< .001** |
| **ACHE** | **0.006 (0.018)** | **-0.083 (0.024)** | **10.05** | **0.022** |
| BACE1 | 0.006 (0.007) | 0.017 (0.008) | 1.42 | 0.236 |
| BASP1 | -0.011 (0.047) | -0.073 (0.060) | 0.95 | 0.331 |
| CD63 | 0.027 (0.025) | 0.029 (0.017) | 0.00 | 0.990 |
| CST3 | 0.011 (0.007) | 0.003 (0.009) | 0.56 | 0.461 |
| **GFAP** | **0.069 (0.019)** | **0.018 (0.018)** | **4.23** | **0.042** |
| IGFBP7 | 0.014 (0.010) | 0.011 (0.008) | 0.21 | 0.649 |
| KLK6 | 0.041 (0.014) | 0.029 (0.010) | 0.78 | 0.378 |
| MAPT | 0.035 (0.024) | 0.039 (0.023) | 0.01 | 0.946 |
| NfL | 0.140 (0.030) | 0.103 (0.025) | 1.05 | 0.307 |
| **PSEN1** | **0.004 (0.014)** | **0.072 (0.023)** | **5.99** | **0.016** |
| pTau-181 | 0.013 (0.023) | 0.069 (0.021) | 3.57 | 0.061 |
| pTau-217 | 0.004 (0.017) | 0.048 (0.016) | 3.61 | 0.060 |
| pTau-231 | 0.025 (0.021) | 0.051 (0.024) | 0.76 | 0.379 |
| SRFP1 | 0.091 (0.050) | 0.142 (0.044) | 0.51 | 0.477 |
| **Aβ42/Aβ40** | **-0.057 (0.014)** | **0.037 (0.022)** | **13.78** | **< .001** |
| **pTau-217/Aβ42** | **0.009 (0.021)** | **-0.057 (0.025)** | **4.22** | **0.042** |

**Bolded** values indicate $p < .05$ , one-tailed test

### Sex Differences and Their Interaction with Age Within Species

We next tested for sex differences and whether the slope in change with age differed between males and females within each species. Analyses of covariance (ANCOVA) were performed on each individual biomarker measure within each species. Sex was the between-group factor while age was the covariate. No significant interactions between sex and age were found for any of the biomarkers within each species. Thus, the patterns of association between age and each biomarker did not differ between sexes within either the rhesus monkey or baboon samples.

### Sex and Species Differences

For these analyses, analysis of covariance was performed on each biomarker with species and sex as between group factors while age was the covariate. Rhesus monkeys had significantly higher NPQ values than the baboons for Aβ38 *F*(1, 130)=6.46, *p* = 0.012, Aβ40 *F*(1, 130) = 11.38 *p* < 0.001, ACHE F(1,133)=55.75, p<0.001, CD63 *F*(1,132) = 4.13, *p* = 0.044, CST3 *F*(1, 132) = 60.52, *p* < 0.001, GFAP F(1,133)=55.93, p<0.001, IFGBP7 *F*(1, 132) = 8.21, *p* = 0.005, pTau-181 *F*(1, 131) = 6.68, *p* < 0.010, pTau-217 *F*(1, 130) = 13.40, *p* < 0.001, and SFRP1 *F*(1, 32) = 27.75, *p* < 0.001 (see Table 3). By contrast, baboons had significantly higher NPQ values than rhesus monkeys for Aβ42 *F*(1, 128) = 12.91, *p* < 0.001, BACE1 *F*(1, 132) = 5.12, *p* = 0.025, and PSEN1 *F*(1, 132) = 16.62, *p* < 0.001. No significant species differences were found for BASP1, KLK6, MAPT, NfL, and pTau-231 (see Table 3). Significant species differences were also found for the three ratio measures including Aβ42/Aβ40 *F*(1, 128) = 49.39, *p* < 0.001, pTau-217/Aβ42 *F*(1, 126) = 32.79, *p* < 0.001 and pTau-217/MAPT *F*(1, 130) = 39.06, *p* < 0.001, with baboons having higher Aβ42/Aβ40 ratios and rhesus monkeys having higher pTau-217/Aβ42 and pTau-217/MAPT ratios (see Table 3). The mean NPQ values for each species and biomarker are shown in Figure 2.

**Figure 2.**
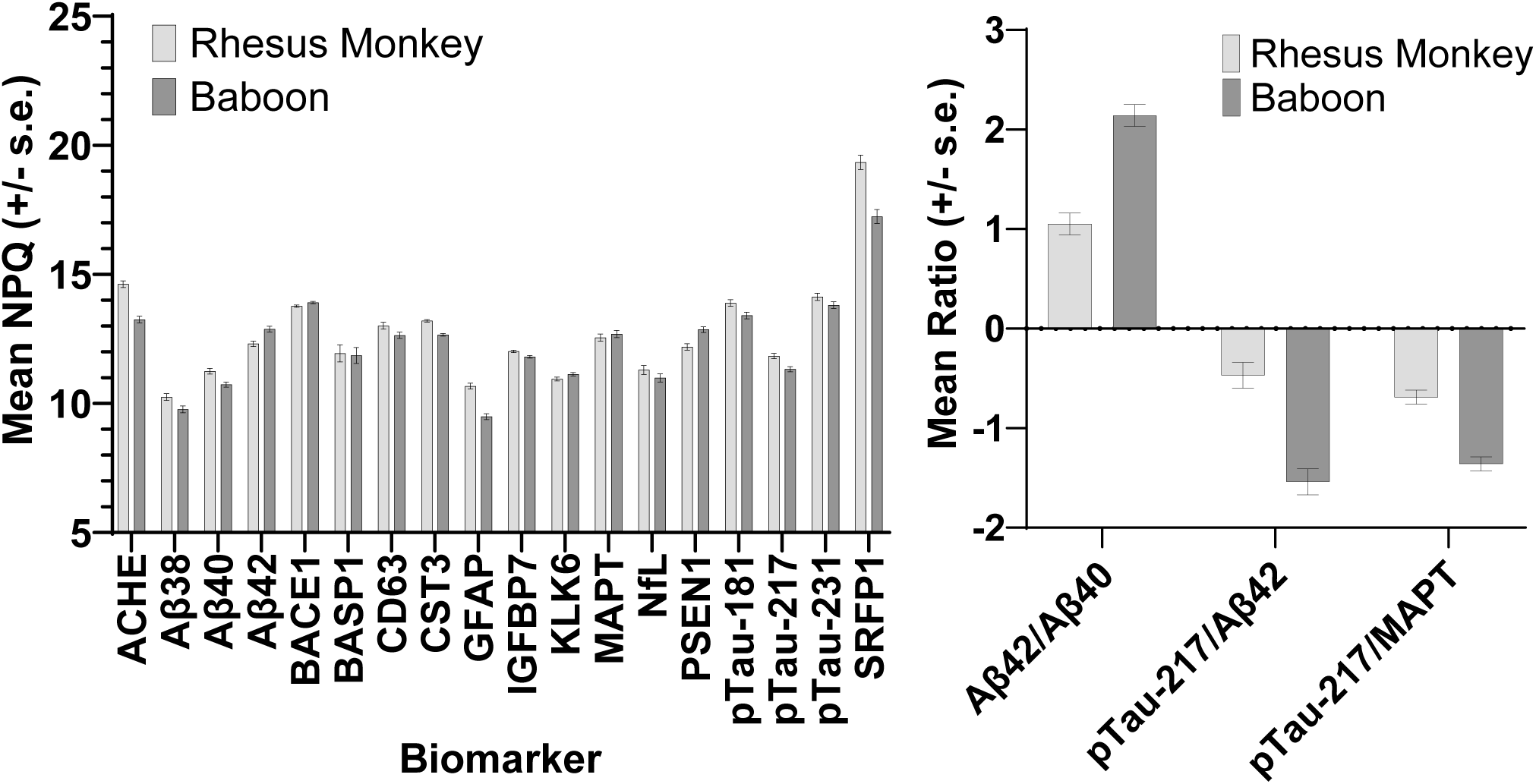
Average rhesus monkey and baboon NPQ values for each biomarker (left) and each ratio measure (right).

**Table 3.** Mean NPQ values (+/- SEM) for Rhesus Monkeys and Baboons for Each Biomarker.

|  | Rhesus<br>M (SEM) | Baboon<br>M (SEM) | Difference |
| --- | --- | --- | --- |
| Aβ38 | 10.25 (0.13) | 9.77 (0.13) | R > B |
| Aβ40 | 11.25 (0.11) | 10.73 (0.10) | R > B |
| Aβ42 | 12.31 (0.11) | 12.88 (0.11) | B > R |
| ACHE | 14.62 (0.13) | 13.25 (0.13) | R > B |
| BACE1 | 13.77 (0.05) | 13.91 (0.04) | B > R |
| BASP1 | 11.94 (0.33) | 11.86 (0.31) | B = R |
| CD63 | 13.01 (0.13) | 12.64 (0.12) | R > B |
| CST3 | 13.2 (0.05) | 12.66 (0.05) | R > B |
| GFAP | 10.67 (0.11) | 9.49 (0.11) | R > B |
| IGFBP7 | 12.02 (0.05) | 11.81 (0.05) | R > B |
| KLK6 | 10.95 (0.07) | 11.13 (0.07) | B = R |
| MAPT | 12.55 (0.14) | 12.69 (0.14) | B = R |
| NfL | 11.3 (0.17) | 10.99 (0.16) | B = R |
| PSEN1 | 12.19 (0.12) | 12.86 (0.11) | B > R |
| pTau-181 | 13.89 (0.13) | 13.41 (0.13) | R > B |
| pTau-217 | 11.84 (0.10) | 11.33 (0.10) | R > B |
| pTau-231 | 14.13 (0.14) | 13.81 (0.13) | R = B |
| SRFP1 | 19.34 (0.28) | 17.24 (0.27) | R > B |
| Aβ42/Aβ40 | 1.05 (0.11) | 2.14 (0.11) | B > R |
| pTau-217/Aβ42 | -0.47 (0.13) | -1.54 (0.13) | R > B |
| pTau-217/MAPT | -0.69 (0.07) | -1.36 (0.07) | R > B |

Values in parentheses are standard error values.

Significant main effects for sex were found for Aβ38 *F*(1, 130)=3.99, *p* = 0.048, Aβ40 *F*(1, 130) = 9.91, *p* < 0.002, Aβ42 *F*(1, 128) = 4.91, *p* = 0.028, CST3 *F*(1, 132) = 15.17, *p* < 0.001, IFGBP7 *F*(1, 132) = 15.34, *p* < 0.001, KLK6 *F*(1, 132) = 8.76, *p* = 0.004, and PSEN1 *F*(1, 132) = 4.08, *p* = 0.046. For each biomarker, males had higher values than females. Finally, a significant two-way interaction was found between species and sex for ACHE *F*(2, 132) = 4.21, *p* = 0.042. Post-hoc analysis indicated that male and female rhesus monkeys did not significantly differ. By contrast, among baboons, males had higher values than females.

### Associations Between Gray Matter Volume and Biomarkers

Within each species, we regressed each biomarker on whole brain gray matter volumes while controlling for sex and age and the results are shown in Table 4. For both rhesus monkeys and baboons, gray matter volume negatively correlated with CST3 values and positively correlated with the Aβ42/Aβ40 ratio (see Figure 3). Additionally, within the baboon sample, gray matter volume significantly negatively correlated with CD63.

**Figure 3.**
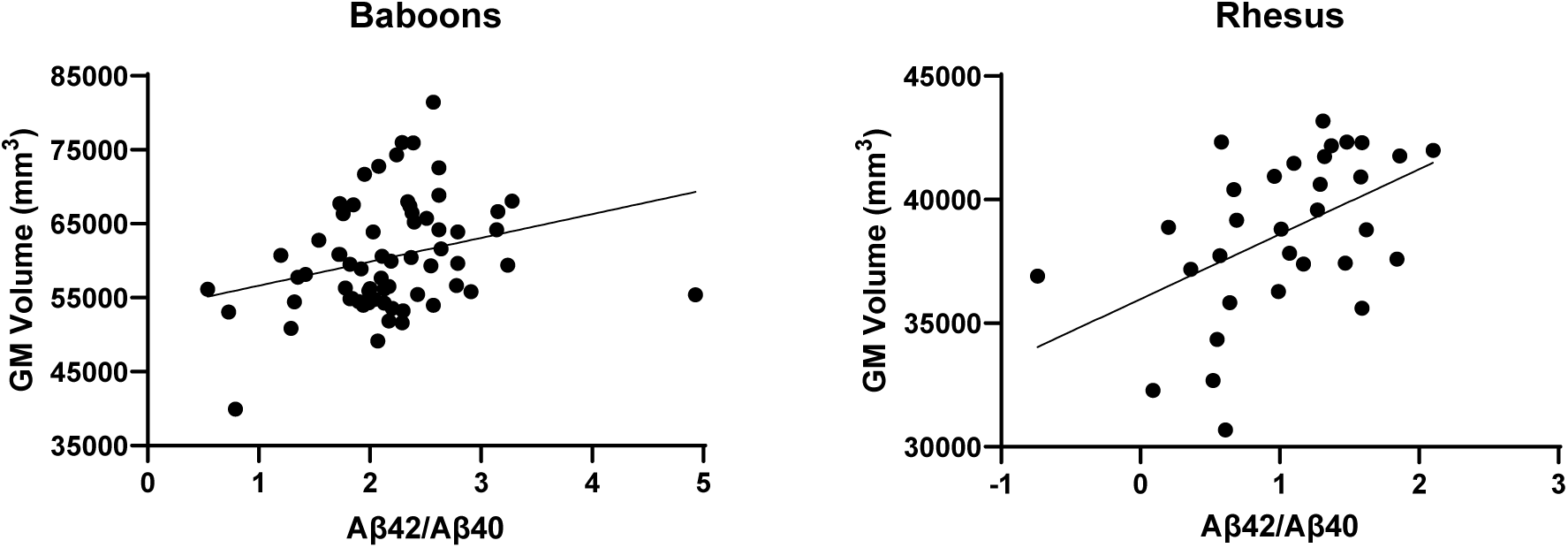
Scatterplots showing the associations between gray matter volume and the plasma Aβ42/Aβ40 ratio for both baboons and rhesus monkeys.

**Table 4.** Partial Correlation Coefficients Between Gray Matter Volume and Each Biomarker Controlling for Sex and Age.

| Biomarker | Rhesus | Baboons |
| --- | --- | --- |
| Aβ38 | -0.06 | -0.07 |
| Aβ40 | -0.313 | -0.184 |
| Aβ42 | 0.121 | 0.081 |
| ACHE | -0.028 | -0.071 |
| BACE1 | -0.03 | 0.029 |
| BASP1 | -0.11 | -0.211 |
| CD63 | 0.027 | -0.289 * |
| CST3 | -0.421 * | -0.282 * |
| GFAP | 0.014 | 0.178 |
| IGFBP7 | -0.124 | -0.007 |
| KLK6 | 0.059 | 0.142 |
| MAPT | 0.028 | 0.093 |
| NfL | -0.263 | -0.01 |
| PSEN1 | 0.072 | 0.08 |
| pTau-181 | -0.008 | 0.17 |
| pTau-217 | 0.065 | 0.222 |
| pTau-231 | 0.035 | 0.123 |
| SRFP1 | -0.256 | -0.079 |
| Aβ42/Aβ40 | 0.397 * | 0.278 ** |
| pTau-217/Aβ42 | -0.012 | 0.008 |

\* $p < .05$ . \*\* $p < .01$

## Discussion

There were three main findings from this study, first, within species, significant associations were found between age and multiple measures of neurodegeneration. Second, between species, significant differences were observed between rhesus monkeys and baboons in several biomarkers, and the patterns of association between age and multiple biomarkers also differed between species. Lastly, within both rhesus monkeys and baboons, we found modest correlations between certain biomarkers and measures of whole-brain gray matter volume.

### Amyloid beta: Aβ38, Aβ40, Aβ42, and Aβ42/Aβ40

We found only partial support for our hypothesis that rhesus monkeys would have higher levels of amyloid biomarkers and stronger associations between amyloid and age compared to baboons. While controlling for age and sex, rhesus monkeys showed significantly higher Aβ38 and Aβ40 NPQ values compared to baboons, consistent with our hypotheses and previous findings in this species (Zhao et al., 2017). However, baboons had significantly higher Aβ42 levels than rhesus monkeys. Moreover, baboons showed a significant quadratic relationship between age and Aβ42 levels, whereas no significant associations with age were found in the rhesus monkeys.

Given the clinical importance of the Aβ42/Aβ40 ratio in predicting amyloid accumulation in the brain and its recommended use for Alzheimer’s disease (AD) diagnosis in humans (Doecke et al., 2020; Jack et al., 2024; Perez-Grijalba et al., 2019; Vergallo et al., 2019), arguably the most interesting finding between the rhesus monkeys and baboons was observed for this measure. For the Aβ42/Aβ40 ratio measure, rhesus monkeys had significantly lower values than baboons, though both species showed significant associations between age, as has been previously reported in rhesus monkeys (Metzler et al., 2025). That said, the pattern of association differed between species. Baboons showed a significant quadratic association with decline, with younger and older baboons exhibiting lower NPQ values than middle-aged individuals, and declines appearing at approximately 15-16 years of age (see Figure 1F). By contrast, rhesus showed a significant negative linear association between age and Aβ42/Aβ40. Thus, older rhesus monkeys and baboons had lower values, which is consistent with reports of human subjects with and without APOE4 mutations (Nakamura et al., 2023; Nakamura et al., 2018). Age-related decline in Aβ42/Aβ40 occurs at an earlier age and appears to be driven by an increase in Aβ40 rather than a decrease in Aβ42 in the rhesus monkeys compared to baboons. Longitudinal data should be collected to examine this pattern further and directly measure changes in these markers as individuals age. As Aβ42/Aβ40 is linked to cognitive declines in humans with disease (Aschenbrenner et al., 2022), future studies should examine the relationship between cognitive function and Aβ42/Aβ40 in rhesus monkeys and baboons.

### Phosphorylated tau: pTau-181, pTau-217, pTau-231 and pTau-217/Aβ42

Consistent with our hypothesis, for pTau-181, pTau-217 and, to a lesser extent, pTau-231, baboons showed a larger slope of change with age compared to rhesus monkeys. These findings are consistent with previously published neuropathology data suggesting that the age of onset of changes in phosphorylated tau occurs earlier in the baboon lifespan compared to rhesus monkeys (Paspalas et al., 2018; Schultz, Hubbard, et al., 2000). The baboons showed significant positive relationships between age and pTau-181, pTau-217 and pTau-231, while the rhesus showed no age-related associations with any of the pTau measures. These findings are consistent with previous data assessing amyloid and tau biomarkers in plasma and CSF in these two species (Metzler et al., 2025; Neal et al., 2025), with the exception of a single report in rhesus monkeys reporting a positive association between age and pTau-217 assayed from CSF (Datta et al., 2024). Similar to the individual phosphorylated tau measures, for the pTau-217/Aβ42 ratio, only baboons showed a significant association with age, with younger and older individuals exhibiting higher NPQ values compared to middle-aged baboons.

### Additional Biomarkers

Cystatin-C (CST3) is a well-studied protein implicated in aging, AD, and other forms of neurodegeneration (Chen et al., 2021; Mathews & Levy, 2016). CST3 has been reported to be co-expressed with amyloid in rhesus monkeys and squirrel monkeys and is a target of some anti-amyloid clinical interventions in patients with probable AD (Walker, 1997; Walker et al., 1990). Here, we found significant positive associations with age in both species and no differences in the mean NPQ values between them. Interestingly, CST3 levels were significantly negatively correlated with Aβ42/Aβ40 ratio measures in both the rhesus monkeys (*r* = -.535, *p* < .001) and baboons (*r* = -.352, *p* = .002); thus, rhesus monkeys and baboons with lower Aβ42/Aβ40 (which potentially reflects increased amyloid pathology in the brain) had higher CST3.

Similar to CST3, we found significant positive associations with age for kallikrein-related peptidase 6 (KLK6) and neurofilament light chain (NfL) in both species, and no differences in the NPQ values between them. KLK6 is a serine protease that is related to brain aging and implicated in AD and other dementias in humans (Ashby et al., 2010; Liu et al., 2025; Patra et al., 2018). CSF and, to a lesser extent, plasma KLK6 levels have been reported to increase with age, and plasma KLK6 levels are significantly higher in individuals with an AD diagnosis compared to controls (Patra et al., 2018). NFL is a biomarker of neuronal damage and cell loss that increases as adult humans age (Khalil et al., 2020). Our findings align with those in humans and in nonhuman primates, including rhesus monkeys and marmosets (Metzler et al., 2025; Phillips et al., 2024).

We also found that PSEN1, SRFP1, and BACE1 increase with age in baboons but not in the rhesus monkeys, and the slope of change differed significantly for PSEN1 between the two species. Individuals with a mutation in the PSEN1 gene are at risk for early onset AD (Kelleher & Shen, 2017), and carriers of the mutation differ in the age of onset of various biomarkers associated with AD in humans (Malotaux et al., 2026). We have no knowledge of previous studies that have examined PSEN1 protein levels in plasma from nonhuman primates; however, recent studies in genetically manipulated cynomolgus monkeys and marmosets have reported significant differences in some plasma biomarkers of neurodegeneration (Li et al., 2024; Sukoff Rizzo et al., 2023). GFAP was also found to significantly increase with age in rhesus monkeys but not in the baboons. GFAP plays a role in astrogliosis and neuroinflammation and has been found to predict rates of cognitive decline in aging humans (Chatterjee et al., 2021). To what extent species differences in GFAP contribute to age-related decline in cognition among nonhuman primates merits further investigation.

### Associations Between Biomarkers and Gray Matter Volume

Finally, despite significant differences in Aβ42/Aβ40 ratio values between species, lower ratios were associated with lower gray matter volumes in both rhesus monkeys and baboons. We believe this is the first evidence demonstrating an association between *in vivo* measures of gray matter and this particular biomarker in nonhuman primates. The Aβ42/Aβ40 ratio is a frequently used and validated biomarker of amyloid pathology, quantified using PET imaging *in vivo* and in postmortem brain tissue (Baiardi et al., 2019; Doecke et al., 2020; Perez-Grijalba et al., 2019; Salvado, Ossenkoppele, et al., 2023; Vergallo et al., 2019). If one assumes that lower gray matter volumes reflect declines in neural integrity (i.e., neuron loss, synapse loss, etc.), then these findings further validate the use of both rhesus monkeys and baboons in studies on mechanisms associated with neurodegeneration, as well as their inclusion in preclinical studies designed to develop and test new treatments for dementias.

### Limitations

Though there are many strengths and a number of unique findings from the data generated in this study, there are also several limitations. First, the sample size for the tests for association between gray matter volume and each biomarker is somewhat small, particularly for the rhesus monkeys. It is important that the current findings be replicated in a larger sample of monkeys. Second, the patterns of association between age and some biomarkers differed between species (see Table 1). While these findings are interesting in and of themselves, they do complicate the analyses and interpretation of the results. Third, the analysis of associations between age and biomarkers were based on cross-sectional data. Like humans, nonhuman primates (including the subjects in this study) are subject to cohort effects. Ideally, to better capture age-related changes in each biomarker, data collected using longitudinal methods should be used. Finally, we analyzed plasma because it is more easily attainable; however, it has some limitations because it contains circulating proteins derived from both peripheral and central nervous systems sources. It will be important in future studies to assess the same biomarkers in CSF as a means of validating and interpreting the data derived from plasma alone.

### Conclusion

In summary, this is the first systematic comparative study of age-related changes in biomarkers of neurodegeneration in two closely related species of nonhuman primates using comparable sample sizes, age ranges, and identical assay procedures. Though many similarities were found in the pattern of aging between species, baboons and rhesus monkeys differed in the relative amounts (or magnitude) and in the time course of changes across multiple biomarkers. For the phosphorylated tau measures, though rhesus monkeys had higher absolute NPQ values, baboons showed significant and stronger associations with age. By contrast, rhesus monkeys appear to exhibit a greater amyloid burden, at least when considering Aβ42/Aβ40 ratio compared to baboons. Collectively, these data suggest that rhesus monkeys and baboons are both important model species for future preclinical studies, with baboons potentially having enhanced translational value for therapeutics designed to target tauopathies.

## Acknowledgements

This work was supported in part by NIH grants AG067419 (WDH), AG087945 (WDH), AG078411 (MMM), AG087914 (MMM), NS128190 (HS), as well as a grant from Cattlemen for Cancer Research (WDH and MMM). The baboon care and collection of biologics were supported in part by the NIH under award number P40OD024628 - SPF Baboon Research Resource. Data were generated in part by the FCCIF-ORION Core, which receive partial support from the National Cancer Institute under grant P30CA016672 to MD Anderson Cancer Center. The research reported here was not directly funded through the P30CA016672 grant to MD Anderson Cancer Center and is not within the scope of such grant.

## Notes

### Competing Interest Statement

The authors have declared no competing interest.

