## Supplemental Table 1 for "A Comparative Study of Plasma Biomarkers of Neurodegeneration in Rhesus Monkeys (*Macaca mulatta*) and Baboons (*Papio anubis*)"

**Supplemental Table 1**. NULISAseq CNS Disease Panel 120 detection rates, and range, mean and standard deviation of NPQ values, and correlations with age for each species (for biomarkers with detection rates over 60%).

| | | Rhesus | | | | | Baboon | | | | |
|---|---|---|---|---|---|---|---|---|---|---|---|
| Biomarker | Detection (%) | Minimum | Maximum | Mean | SD | r (age) | Minimum | Maximum | Mean | SD | r (age) |
| ACHE | 100 | 12.92 | 15.81 | 14.44 | 0.60 | 0.045 | 12.29 | 15.98 | 13.43 | 0.80 | **-0.376** |
| AGRN | 100 | 10.42 | 12.00 | 11.22 | 0.32 | 0.197 | 10.50 | 11.64 | 11.03 | 0.28 | 0.028 |
| ANXA5 | 100 | 10.39 | 20.42 | 16.24 | 2.36 | **0.253** | 10.76 | 20.72 | 17.90 | 2.36 | **0.418** |
| APOE | 14 | | | | | | | | | | |
| APOE4 | 11.6 | | | | | | | | | | |
| ARSA | 100 | 6.71 | 12.72 | 9.71 | 1.46 | 0.041 | 7.08 | 14.53 | 11.24 | 1.35 | **0.232** |
| Aβ38 | 98.8 | 8.45 | 12.10 | 10.16 | 0.63 | **0.4** | 6.51 | 11.24 | 9.61 | 0.89 | **0.374** |
| Aβ40 | 98.8 | 9.35 | 12.23 | 11.15 | 0.59 | **0.416** | 8.94 | 11.91 | 10.50 | 0.70 | **0.362** |
| Aβ42 | 96.5 | 11.49 | 13.30 | 12.27 | 0.42 | -0.04 | 10.27 | 14.91 | 12.68 | 0.78 | **0.463** |
| BACE1 | 100 | 13.19 | 14.15 | 13.72 | 0.25 | 0.103 | 13.26 | 14.31 | 13.81 | 0.25 | **0.26** |
| BASP1 | 90.7 | 9.85 | 16.59 | 11.59 | 1.60 | -0.031 | 10.22 | 18.90 | 12.10 | 1.85 | -0.143 |
| BDNF | 67.4 | 4.07 | 14.99 | 9.33 | 2.13 | 0.119 | 1.44 | 11.10 | 5.43 | 1.71 | 0.131 |
| CALB2 | 84.9 | 7.32 | 11.40 | 8.76 | 0.68 | -0.184 | 8.08 | 12.82 | 9.27 | 0.71 | -0.055 |
| CCL11 | 37.2 | | | | | | | | | | |
| CCL13 | 100 | 6.04 | 10.29 | 8.37 | 0.98 | -0.08 | 5.89 | 11.39 | 8.86 | 1.18 | 0.188 |
| CCL17 | 27.7 | | | | | | | | | | |
| CCL2 | 100 | 7.97 | 10.38 | 9.23 | 0.61 | **0.283** | 7.73 | 12.48 | 9.21 | 0.79 | -0.039 |
| CCL22 | 100 | 6.36 | 10.63 | 8.05 | 0.90 | -0.135 | 5.89 | 9.33 | 7.67 | 0.78 | 0.061 |
| CCL26 | 97.7 | 4.79 | 11.57 | 6.31 | 1.10 | -0.011 | 5.94 | 12.46 | 7.58 | 0.92 | -0.064 |
| CCL3 | 90.7 | 1.34 | 6.76 | 3.76 | 1.35 | 0.292 | 3.04 | 9.00 | 6.02 | 1.52 | 0.131 |
| CCL4 | 93 | 4.32 | 14.50 | 7.66 | 1.36 | 0.082 | 3.03 | 9.24 | 6.06 | 0.95 | -0.008 |
| CD40LG | 100 | 7.79 | 17.95 | 13.00 | 1.98 | **0.351** | 10.86 | 19.46 | 15.73 | 1.94 | **0.41** |
| CD63 | 100 | 11.80 | 15.76 | 12.91 | 0.85 | 0.142 | 11.26 | 13.90 | 12.53 | 0.55 | 0.18 |
| CHI3L1 | 100 | 12.16 | 18.38 | 15.39 | 1.26 | **0.257** | 11.92 | 16.26 | 13.49 | 0.85 | 0.032 |
| CHIT1 | 100 | 9.81 | 16.48 | 12.91 | 1.16 | **0.27** | 12.29 | 19.61 | 14.16 | 1.02 | 0.063 |
| CNTN2 | 2.3 | | | | | | | | | | |
| CRH | 90.7 | 8.90 | 26.03 | 13.98 | 5.51 | -0.063 | 9.02 | 29.06 | 14.03 | 6.29 | **-0.292** |
| CRP | 100 | 11.19 | 17.26 | 13.99 | 1.25 | **-0.441** | 15.09 | 17.61 | 15.69 | 0.48 | 0.034 |
| CSF2 | 100 | 9.94 | 15.60 | 13.29 | 1.20 | 0.111 | 10.89 | 14.28 | 12.61 | 0.67 | -0.047 |

| | | | | | | | | | | | |
|---|---|---|---|---|---|---|---|---|---|---|---|
| CST3 | 100 | 12.28 | 13.63 | 13.11 | 0.27 | 0.192 | 11.87 | 13.07 | 12.56 | 0.28 | 0.04 |
| CX3CL1 | 18.6 | | | | | | | | | | |
| CXCL1 | 0 | | | | | | | | | | |
| CXCL10 | 100 | 10.28 | 13.63 | 11.48 | 0.71 | -0.057 | 9.45 | 17.44 | 11.62 | 1.12 | -0.091 |
| CXCL8 | 98.8 | 7.41 | 18.01 | 11.35 | 2.66 | **0.373** | 13.35 | 21.75 | 18.72 | 1.73 | **0.37** |
| DDC | 100 | 13.65 | 15.68 | 14.87 | 0.48 | -0.092 | 12.29 | 15.72 | 14.22 | 0.68 | -0.187 |
| ENO2 | 2.3 | | | | | | | | | | |
| FABP3 | 100 | 7.91 | 15.93 | 11.21 | 1.79 | 0.245 | 5.32 | 13.56 | 10.01 | 1.78 | -0.056 |
| FCN2 | 3.5 | | | | | | | | | | |
| FGF2 | 100 | 11.04 | 18.02 | 14.91 | 1.58 | **0.308** | 10.75 | 17.21 | 14.46 | 1.57 | **0.288** |
| FLT1 | 100 | 13.73 | 22.19 | 16.23 | 1.95 | 0.035 | 13.65 | 21.82 | 15.62 | 2.03 | **-0.383** |
| FOLR1 | 73.3 | 0.00 | 8.14 | 5.43 | 1.32 | -0.032 | 0.00 | 7.40 | 4.25 | 1.83 | -0.011 |
| GDF15 | 100 | 11.07 | 18.96 | 13.28 | 2.07 | -0.001 | 12.35 | 18.92 | 14.06 | 1.80 | **-0.273** |
| GDI1 | 100 | 8.24 | 11.06 | 9.45 | 0.62 | -0.195 | 9.33 | 13.44 | 10.87 | 0.84 | **0.407** |
| GDNF | 65.1 | 9.01 | 17.03 | 10.75 | 1.83 | -0.084 | 8.99 | 19.48 | 10.62 | 1.68 | 0.142 |
| GFAP | 87.2 | 9.07 | 13.54 | 10.60 | 0.72 | **0.43** | 7.97 | 11.06 | 9.39 | 0.56 | 0.115 |
| GOT1 | 100 | 10.58 | 14.00 | 11.84 | 0.59 | -0.086 | 10.87 | 13.19 | 11.88 | 0.48 | 0.182 |
| HBA1 | 29.1 | | | | | | | | | | |
| HTT | 100 | 11.22 | 16.73 | 14.25 | 1.40 | 0.14 | 10.94 | 17.58 | 14.58 | 1.41 | **0.396** |
| ICAM1 | 1.2 | | | | | | | | | | |
| IFNG | 2.3 | | | | | | | | | | |
| IGF1R | 100 | 12.18 | 13.56 | 12.71 | 0.29 | 0.101 | 12.61 | 14.14 | 13.23 | 0.29 | -0.21 |
| IGFBP7 | 100 | 11.08 | 12.90 | 11.90 | 0.38 | 0.177 | 11.07 | 12.23 | 11.73 | 0.25 | 0.16 |
| IL10 | 8.1 | | | | | | | | | | |
| IL12p70 | 0 | | | | | | | | | | |
| IL13 | 0 | | | | | | | | | | |
| IL15 | 100 | 12.42 | 13.90 | 13.11 | 0.34 | -0.051 | 11.17 | 13.20 | 12.14 | 0.37 | **-0.272** |
| IL16 | 100 | 8.56 | 11.37 | 9.89 | 0.60 | -0.107 | 9.29 | 12.89 | 10.51 | 0.70 | 0.08 |
| IL17A | 87.2 | 8.95 | 22.22 | 12.09 | 1.99 | 0.063 | 9.35 | 15.50 | 11.93 | 1.26 | 0.17 |
| IL18 | 100 | 11.33 | 13.95 | 12.52 | 0.50 | 0.119 | 11.34 | 15.72 | 12.75 | 0.74 | -0.058 |
| IL1B | 4.7 | | | | | | | | | | |
| IL2 | 1.2 | | | | | | | | | | |
| IL33 | 100 | 13.42 | 16.69 | 14.80 | 0.69 | -0.204 | 13.13 | 17.08 | 14.87 | 0.78 | -0.026 |
| IL4 | 16.3 | | | | | | | | | | |

| | | | | | | | | | | | |
|---|---|---|---|---|---|---|---|---|---|---|---|
| IL5 | 100 | 8.56 | 17.87 | 11.62 | 1.35 | **0.266** | 8.30 | 15.40 | 10.99 | 1.29 | -0.023 |
| IL6 | 100 | 9.35 | 16.34 | 12.16 | 1.34 | 0.248 | 11.87 | 15.40 | 13.48 | 0.67 | 0.089 |
| IL6R | 87.2 | 7.37 | 10.19 | 8.87 | 0.58 | 0.195 | 6.64 | 9.74 | 8.28 | 0.65 | 0.073 |
| IL7 | 100 | 10.03 | 17.78 | 13.67 | 1.64 | **0.283** | 9.74 | 15.08 | 12.28 | 1.31 | **0.38** |
| IL9 | 0 | | | | | | | | | | |
| KDR | 8.1 | | | | | | | | | | |
| KLK6 | 100 | 9.58 | 12.35 | 10.83 | 0.54 | **0.354** | 10.38 | 11.90 | 11.04 | 0.33 | **0.314** |
| MAPT | 100 | 11.05 | 15.79 | 12.50 | 0.83 | 0.184 | 11.38 | 15.06 | 12.63 | 0.73 | 0.194 |
| MDH1 | 0 | | | | | | | | | | |
| MME | 100 | 3.08 | 10.96 | 6.72 | 1.63 | 0.05 | 0.00 | 8.70 | 3.62 | 1.46 | -0.212 |
| MSLN | 4.7 | | | | | | | | | | |
| NEFH | 100 | 12.75 | 25.58 | 19.44 | 2.30 | **0.469** | 13.60 | 23.12 | 19.96 | 2.31 | **0.431** |
| NEFL | 96.5 | 9.28 | 14.86 | 11.40 | 1.19 | **0.517** | 9.07 | 13.60 | 10.90 | 0.85 | **0.436** |
| NGF | 100 | 7.72 | 17.09 | 10.10 | 2.32 | 0.007 | 7.15 | 17.90 | 10.66 | 2.81 | **-0.281** |
| NPTX1 | 96.5 | 9.82 | 15.31 | 11.37 | 0.82 | 0.238 | 9.64 | 12.02 | 11.02 | 0.48 | 0.167 |
| NPTX2 | 100 | 10.12 | 13.05 | 11.08 | 0.64 | 0.202 | 11.45 | 14.75 | 13.05 | 0.69 | **0.269** |
| NPTXR | 100 | 8.32 | 11.54 | 9.95 | 0.61 | 0.187 | 9.02 | 11.69 | 10.54 | 0.49 | -0.051 |
| NPY | 98.8 | 4.86 | 12.45 | 6.89 | 1.27 | 0.028 | 3.01 | 11.23 | 7.12 | 1.26 | 0.147 |
| NRGN | 100 | 6.90 | 16.33 | 12.39 | 2.04 | **0.356** | 9.39 | 19.89 | 15.88 | 2.26 | **0.458** |
| Oligo-SNCA | 96.5 | 6.77 | 20.74 | 11.28 | 3.14 | -0.217 | 7.40 | 21.81 | 12.38 | 3.66 | 0.208 |
| PARK7 | 96.5 | 8.74 | 18.28 | 11.17 | 1.41 | -0.022 | 8.24 | 14.94 | 12.50 | 1.04 | 0.195 |
| PDGFRB | 100 | 11.30 | 13.02 | 12.16 | 0.39 | 0.186 | 12.17 | 14.49 | 13.56 | 0.50 | 0.137 |
| PDLIM5 | 86 | 7.84 | 13.74 | 10.06 | 1.73 | 0.087 | 8.39 | 20.36 | 10.37 | 1.85 | 0.023 |
| PGF | 96.5 | 10.05 | 13.20 | 11.98 | 0.65 | -0.005 | 10.10 | 12.36 | 11.61 | 0.55 | **0.363** |
| PGK1 | 100 | 6.08 | 16.43 | 10.77 | 1.91 | -0.083 | 7.60 | 16.56 | 12.94 | 2.04 | **0.441** |
| POSTN | 100 | 6.46 | 13.64 | 10.32 | 1.37 | 0.021 | 8.56 | 13.99 | 12.25 | 1.08 | 0.154 |
| PRDX6 | 0 | | | | | | | | | | |
| PSEN1 | 100 | 10.73 | 13.08 | 12.10 | 0.49 | 0.042 | 10.86 | 14.84 | 12.72 | 0.75 | **0.345** |
| pSNCA-129 | 100 | 10.95 | 18.04 | 13.31 | 1.71 | **-0.265** | 11.25 | 18.62 | 13.56 | 1.85 | **0.232** |
| pTau-181 | 100 | 12.56 | 16.64 | 13.90 | 0.76 | 0.074 | 11.59 | 14.87 | 13.25 | 0.68 | **0.368** |
| pTau-217 | 97.7 | 10.69 | 13.76 | 11.83 | 0.57 | 0.027 | 10.10 | 12.53 | 11.25 | 0.51 | **0.338** |
| pTau-231 | 100 | 12.82 | 16.71 | 14.15 | 0.72 | 0.153 | 11.98 | 16.08 | 13.64 | 0.78 | **0.241** |
| pTDP43-409 | 100 | 8.99 | 15.41 | 11.21 | 1.11 | -0.011 | 10.13 | 14.99 | 12.47 | 0.97 | **0.273** |
| PTN | 14 | | | | | | | | | | |

| | | | | | | | | | | | |
|---|---|---|---|---|---|---|---|---|---|---|---|
| REST | 0 | | | | | | | | | | |
| RUVBL2 | 100 | 9.87 | 15.06 | 12.29 | 1.12 | 0.21 | 10.61 | 16.38 | 13.62 | 1.24 | **0.294** |
| S100A12 | 44.2 | | | | | | | | | | |
| S100B | 100 | 10.74 | 18.03 | 12.73 | 1.52 | -0.112 | 10.81 | 16.75 | 12.86 | 1.01 | 0.031 |
| SAA1 | 0 | | | | | | | | | | |
| SFRP1 | 100 | 16.72 | 25.37 | 19.42 | 1.72 | 0.231 | 13.17 | 22.11 | 17.20 | 1.45 | **0.354** |
| SFTPD | 100 | 12.15 | 16.18 | 14.24 | 0.76 | -0.096 | 11.99 | 14.93 | 13.58 | 0.66 | 0.047 |
| SLIT2 | 100 | 11.61 | 13.43 | 12.43 | 0.39 | 0.223 | 12.85 | 14.21 | 13.33 | 0.27 | -0.047 |
| SMOC1 | 4.7 | | | | | | | | | | |
| SNAP25 | 5.8 | | | | | | | | | | |
| SNCA | 100 | 11.57 | 17.12 | 13.73 | 1.33 | **-0.253** | 12.34 | 17.65 | 14.46 | 1.31 | 0.228 |
| SNCB | 17.4 | | | | | | | | | | |
| SOD1 | 100 | 5.58 | 17.62 | 9.85 | 2.20 | **-0.261** | 6.00 | 15.83 | 10.05 | 2.06 | **0.326** |
| SQSTM1 | 100 | 11.19 | 16.18 | 13.31 | 0.99 | -0.041 | 12.05 | 16.90 | 14.22 | 1.14 | **0.305** |
| TAFA5 | 100 | 5.38 | 9.89 | 7.80 | 0.92 | 0.048 | 4.09 | 13.17 | 8.17 | 1.74 | **0.282** |
| TARDBP | 100 | 11.64 | 18.15 | 15.22 | 1.35 | 0.157 | 12.58 | 18.10 | 16.07 | 1.51 | **0.368** |
| TEK | 14 | | | | | | | | | | |
| TIMP3 | 100 | 12.05 | 21.69 | 17.98 | 2.31 | 0.176 | 12.13 | 21.50 | 18.34 | 1.96 | **0.456** |
| TNF | 5.8 | | | | | | | | | | |
| TREM1 | 0 | | | | | | | | | | |
| TREM2 | 16.3 | | | | | | | | | | |
| UBB | 88.4 | 10.94 | 21.57 | 13.07 | 1.85 | 0.17 | 11.09 | 17.85 | 12.83 | 1.62 | 0.153 |
| UCHL1 | 0 | | | | | | | | | | |
| VCAM1 | 2.3 | | | | | | | | | | |
| VEGFA | 100 | 12.02 | 14.74 | 13.02 | 0.51 | 0.194 | 12.38 | 14.18 | 12.94 | 0.32 | -0.152 |
| VEGFD | 87.2 | 9.77 | 11.57 | 10.56 | 0.38 | -0.166 | 10.05 | 12.39 | 11.16 | 0.49 | **0.282** |
| VGF | 70.9 | 8.95 | 19.56 | 12.48 | 2.42 | -0.151 | 9.12 | 16.81 | 11.21 | 1.33 | **0.329** |
| VSNL1 | 100 | 9.74 | 11.33 | 10.62 | 0.32 | 0.116 | 10.42 | 12.18 | 11.12 | 0.40 | -0.085 |
| YWHAG | 54.7 | | | | | | | | | | |
| YWHAZ | 46.5 | | | | | | | | | | |

**Bold** r values indicate significant correlations between biomarker and age.
